# Deep sequencing of circulating exosomal microRNA allows non-invasive glioblastoma diagnosis

**DOI:** 10.1101/342154

**Authors:** Saeideh Ebrahimkhani, Fatemeh Vafaee, Susannah Hallal, Heng Wei, Maggie Yuk T. Lee, Paul E. Young, Laveniya Satgunaseelan, Brindha Shivalingam, Catherine M. Suter, Michael E. Buckland, Kimberley L. Kaufman

## Abstract

Exosomes are nano-sized extracellular vesicles released by many cells that contain molecules characteristic of their cell-of-origin, including microRNA. Exosomes released by glioblastoma cross the blood-brain-barrier into the peripheral circulation, and carry molecular cargo distinct to that of ‘free-circulating’ miRNA. In this pilot study, serum exosomal-microRNAs were isolated from glioblastoma (*n*=12) patients and analyzed using unbiased deep sequencing. Results were compared to sera from age- and gender-matched healthy controls, and to grades II-III (*n*=10) glioma patients. Significant differentially expressed microRNAs were identified, and the predictive power of individual and subsets of microRNAs were tested using univariate and multivariate analyses. Additional sera from glioblastoma patients (*n*=4) and independent sets of healthy (*n*=9) and non-glioma (*n*=10) controls were used to further test the specificity and predictive power of this unique exosomal-microRNA signature. Twenty-six microRNAs were differentially expressed in serum exosomes from glioblastoma patients’ relative to healthy controls. Random forest modeling and data partitioning selected seven miRNAs (miR-182-5p, miR-328-3p, miR-339-5p, miR-340-5p, miR-485-3p, miR-486-5p and miR-543) as the most stable for classifying glioblastoma. Strikingly, within this model, six iterations of these miRNA classifiers could distinguish glioblastoma patients from controls with perfect accuracy. The seven-miRNA panel was able to correctly classify all specimens in validation cohorts (*n*=23). Also identified were 23 dysregulated miRNAs in IDH^MUT^ gliomas, a partially overlapping yet distinct signature of lower grade glioma. Serum exosomal-miRNA signatures can accurately diagnose glioblastoma preoperatively. miRNA signatures identified are distinct from previously reported ‘free-circulating’ miRNA studies in GBM patients, and appear to be superior.

## INTRODUCTION

Malignant gliomas, particularly glioblastoma (GBM), represent the most lethal primary brain tumors, owing in part to their highly infiltrative growth patterns. The World Health Organization (WHO) guidelines sub-categorize glioma by histopathologic evaluation into tumor grades I-IV, where GBM (grade IV) is the most aggressive and also the most common. Despite surgery, radiation, and chemotherapy, essentially all GBM tumors recur, at which point patients have reduced treatment options and worsening prognoses. Compounding this aggressive cancer phenotype are challenges in monitoring responses to treatment and tumor progression. While recent revisions to the Response Assessment in Neuro-Oncology (RANO) criteria helps to standardize glioma tumor monitoring^1^, radiographic measurements can be unreliable and insensitive to early signs of treatment failure and tumor relapse. Moreover, there are still difficulties deciphering pseudo-progression and pseudo-responses in some patients. Brain biopsy and histologic analysis can provide definitive diagnoses and evaluation of disease progression, however serial biopsies are impractical given the cumulative surgical risk, and biopsied tissue may not reflect the heterogeneity of GBM tumors.

An important step towards the provision of personalized GBM patient care is the ability to assess tumors *in-situ*. As such, there is a real need for biomarkers that can measure disease burden and treatment responses in GBM patients in a safe, accurate and timely manner, and preferably before changes become clinically apparent. The recently popularized idea of ‘liquid biopsy’ presents an ideal approach to monitor GBM tumor load and evolution in response to treatment. If developed and implemented alongside new treatments, such tests would provide useful surrogate endpoints and allow clinical trial protocols to be more dynamic and adaptive.

Exosomes are nano-sized (30-100 nm) membrane-bound extracellular vesicles released by all cells in both health and disease, and there is growing interest in their use as non-invasive biomarkers for disease diagnosis and monitoring of disease recurrence^2^. GBM-derived exosomes circulate in the peripheral blood of patients, and can contain diagnostic nucleic acid ^3^. We recently described a GBM exosome protein signature^4^ and also showed that GBM exosomes contain abundant, selectively packaged small non-coding RNAs (sncRNAs)^5^. Using unbiased sncRNA deep sequencing, we identified several unusual and/or completely novel sncRNAs within GBM exosomes *in vitro* as well as an enrichment of microRNA (miRNA) implicated in oncogenesis, including miR-23a, miR-30a, miR-221 and miR-451^5^. Thus, while GBM exosomal miRNA contents broadly reflect their cell of origin, there is a unique profile of miRNAs within exosomes.

Some studies of exosomal miRNA in GBM patients have already been reported; these studies utilized methods that focused on pre-defined and relatively small groups of miRNA species. One previous study found that miR-21 levels in CSF exosomes of GBM patients were up-regulated 10-fold compared to controls^6^, while another reported that serum exosomal miR-320, mir-547-3p, and RNU6-1 were significantly associated with GBM diagnosis, as well as outcome (RNU6-1)^7^. However, to date no comprehensive analysis of the entire miRNA repertoire of serum exosomes in glioma patients has been performed. Here, we have used unbiased next generation sequencing and an integrative bioinformatics pipeline^8^ to assay the complete repertoire of exosomal-associated miRNAs in the serum of patients with glioblastoma, lower grade gliomas, and healthy controls. We describe a novel miRNA signature within serum exosomes that is highly predictive of pre-operative GBM diagnosis. Furthermore, we show that this approach has potential for describing unique miRNA signatures for distinct glioma entities.

## RESULTS

### Characterization of serum exosomes isolated prior to miRNA sequencing

Serum exosomes were isolated by size exclusion chromatography. The combined elution fractions 8-10 showed particle sizes with a mean diameter 89.1 ± 2.5 nm and modal diameter of 81.7±5.5 nm (**Fig. 1a**). TEM confirmed the presence of similarly sized particles with vesicular morphologies, characteristic of exosomes (**Fig. 1b**). MS analysis confidently identified 1167, 861 and 636 proteins in qEV elution fractions 8, 9 and 10 from healthy serum, respectively (**Supp.Table 2**). Overall, 87 of the top 100 proteins commonly identified in exosomes were confidently sequenced across the three fractions, including all top 10 exosomal proteins (**Fig. 1c-1**). Primary sub-cellular localizations included significant enrichments of ‘exosome’ and ‘blood microparticle’ related proteins across all fractions, with minimal contamination from other compartments, including the nucleolus (**Fig. 1c-2**) where certain miRNAs show specific nuclear enrichment^9^. Prior to RNA extraction, serums were treated with RNaseA to remove circulating RNAs that may confound measurements of exosomal RNAs^8^. RNA extracted from each sample yielded profiles typical for exosomes, showing an absence of ribosomal RNA and enrichment of small (<200 nt) RNA species (**Fig. 1d**).

**Table 1B:** Additional patients and cohorts used for validation

| Patient/cohort | Age | Gender | Diagnosis | Notes |
| --- | --- | --- | --- | --- |
| GBM1_relapse | 46 | M | GBM IV | Pre-operative blood taken after recurrence of GBM1 (8-month relapse) |
| GBM12_prior | 45 | F | GBM IV | Pre-operative blood taken before removal of earlier GBM lesion (GBM12; 4.6 months prior) |
| GBM13 | 33 | M | GBM IV | Glioblastoma, IDH<sup>MUT</sup>, WHO (2016) grade IV |
| GBM14 | 56 | M | high-grade glioma | No surgery/tissue pathology performed, diagnosis based on repeat MRIs. Overall survival of 8.1 months. |
| GI_C | 24 | F | Ganglioglioma grade I | GFAP<sup>+</sup> in glial component/ NeuN<sup>+</sup> in neuronal component, IDH1<sup>WT</sup>, ATRX<sup>+</sup>, BRAF(V600E)<sup>+++</sup> |
| HC ($n=9$) | 36.2 ± 10.3 | 5F, 4M | Healthy controls | - |
| MS ($n=9$) | 35.3 ± 10.4 | 5M, 4F | Relapse-remitting Multiple Sclerosis | All patients had active lesions; were untreated ($n=5$) or receiving different immunomodulatory therapies ($n=4$). |

**Figure 1.**
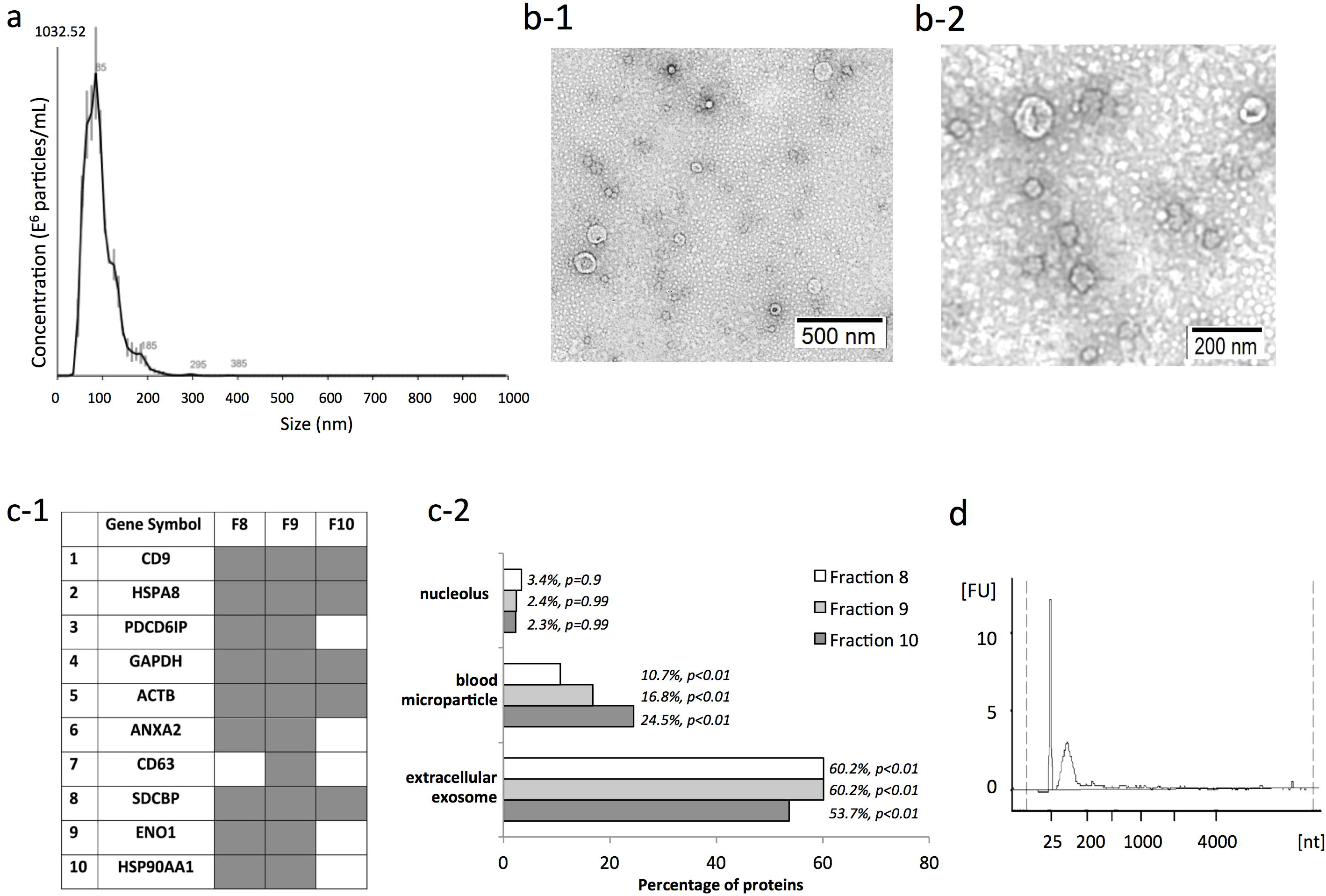
Characterization of serum exosomes isolated in fractions 8-10 by size exclusion chromatography prior to miRNA sequencing. **(a.)** Size distribution of particles as analyzed by nanoparticle tracking analysis. (**b.**) Transmission electron microscopy allowed visualization of vesicles with sizes ranging from 60-110 nm in diameter, scale bars = 500 nm (**b-1.,** *wide field*) and 200 nm (**b-2**,. *close-up*). **(c-1.**) Mass spectrometry-based proteome analysis of size chromatographic elution fractions 8-10 identified all top 10 exosome marker proteins and (**c-2.**) showed significant enrichment of proteins characteristic of exosomes and blood microparticles. Proteins identified in fractions 8-10 showed limited, non-significant associations to compartments like the nucleolus, where certain miRNA species are concentrated. (**d.**) Bioanalyzer trace of RNA extracted from serum exosomes shows the main population of small RNA and no ribosomal RNA.

### Differentially expressed exosomal miRNAs in GBM patient sera

Circulating exosomal miRNA profiles from patients with histopathologically confirmed IDH^WT^ GBM (*n*=12) were compared to age- and gender-matched healthy controls (*n*=12; see **Table 1A** for discovery cohorts; **Table 1B** for validation cases). We employed three statistical approaches (Student’s *t*-test, Fisher’s exact, Wilcoxon rank sum) to identify a discovery set of differentially expressed miRNA biomarkers. miRNA biomarkers were identified if their differential expression met a fold change≥2 in either direction and unadjusted *p*-values≤0.05 in all statistical tests applied. Using this approach, we identified 26 miRNAs significantly dysregulated between healthy controls and GBM patients (**Table 2; Fig. 2-a**; normalized miRNA counts are available in **Supp.Table 3** and differential expression analysis in **Supp.Table 4A**).

**Table 1A:** Overview of cohorts used for discovery miRNA analyses.

|  | GBM, IDH<sup>WT</sup> | GBM-matched HC | GII-III, IDH<sup>MUT</sup> | GII-III-matched HC |
| --- | --- | --- | --- | --- |
| Sample $n$ | 12 | 12 | 10 | 10 |
| Age (mean ± SD) | 63.3 ± 11.5 | 56.2 ± 12.4 | 42.9 ± 12.7 | 42.7 ± 10.2 |
| Gender | 7M, 5F | 7M, 5F | 6M, 4F | 6M, 4F |

**Table 2.** Significant dysregulated miRNAs in serum exosomes from glioblastoma (GBM) patients (*n*=12) relative to healthy controls (HC; *n*=12).

| miRNA | CPM<br>(GBM) | CPM<br>(HC) | FC | Exact<br>test | *t*-test | Wilcoxon | Error<br>rate | AUROC | 95% CI of<br>AUROC |
| --- | --- | --- | --- | --- | --- | --- | --- | --- | --- |
| **486-5p** | 25291.6 | 8522.6 | 3.0 | 1.6E-07* | 4.0E-04* | 1.0E-04* | 0.149 | 0.924 | (0.823, 1) |
| **182-5p** | 2090.5 | 850.6 | 2.5 | 5.7E-07* | 3.0E-04* | 2.0E-04* | 0.151 | 0.917 | (0.808, 1) |
| **486-3p** | 277.4 | 114 | 2.4 | 5.0E-06* | 0.002* | 3.0E-04* | 0.149 | 0.910 | (0.791, 1) |
| **378a-3p** | 2083.2 | 875.2 | 2.4 | 1.4E-06* | 0.003* | 4.0E-04* | 0.158 | 0.903 | (0.783, 1) |
| **183-5p** | 645.8 | 267.9 | 2.4 | 2.0E-05* | 0.001* | 0.001* | 0.176 | 0.882 | (0.749, 1) |
| **501-3p** | 359.6 | 157.3 | 2.3 | 1.1E-05* | 0.002* | 0.001* | 0.161 | 0.875 | (0.726, 1) |
| **20b-5p** | 594.6 | 266.3 | 2.2 | 2.9E-06* | 0.002* | 1.0E-04* | 0.133 | 0.938 | (0.834, 1) |
| **106b-3p** | 2703.2 | 1215 | 2.2 | 3.9E-06* | 0.001* | 0.001* | 0.160 | 0.889 | (0.752, 1) |
| **629-5p** | 896.8 | 415 | 2.2 | 0.001* | 0.047 | 0.04 | 0.235 | 0.743 | (0.532, 0.954) |
| **185-5p** | 23250.5 | 11424.1 | 2.0 | 4.3E-05* | 0.007* | 0.005* | 0.207 | 0.833 | (0.670, 0.997) |
| **25-3p** | 21838.8 | 10949.9 | 2.0 | 0.001* | 0.002* | 0.006* | 0.199 | 0.826 | (0.662, 0.991) |
| **21-5p** | 73535.3 | 142796.9 | -2.0 | 2.7E-04* | 4.2E-05* | 5.0E-05* | 0.133 | 0.944 | (0.862, 1) |
| **7a-3p** | 82.1 | 176.3 | -2.0 | 0.003* | 0.005* | 0.010* | 0.187 | 0.806 | (0.611, 1) |
| **381-3p** | 190.5 | 397.9 | -2.0 | 0.009* | 0.012 | 0.012 | 0.220 | 0.799 | (0.620, 0.977) |
| **409-3p** | 1146.9 | 2242.5 | -2.0 | 0.019 | 0.029 | 0.024 | 0.233 | 0.771 | (0.575, 0.967) |
| **7d-3p** | 1050.5 | 1912.9 | -2.0 | 0.005* | 0.013 | 0.017 | 0.209 | 0.785 | (0.574, 0.996) |
| **323b-3p** | 117.3 | 288.3 | -2.4 | 0.004* | 0.010* | 0.004* | 0.199 | 0.840 | (0.665, 1) |
| **328-3p** | 382.5 | 922.5 | -2.5 | 4.6E-06* | 2.0E-04* | 2.2E-05* | 0.117 | 0.958 | (0.889, 1) |
| **339-5p** | 90.1 | 234.8 | -2.5 | 1.2E-06* | 2.0E-04* | 3.3E-05* | 0.109 | 0.951 | (0.864, 1) |
| **340-5p** | 1536 | 3848.1 | -2.5 | 4.8E-06* | 1.0E-04* | 5.0E-05* | 0.134 | 0.944 | (0.858, 1) |
| **126-5p** | 1222.3 | 2947 | -2.5 | 5.6E-06* | 0.002* | 0.001* | 0.150 | 0.896 | (0.767, 1) |
| **130b-5p** | 111.9 | 248.9 | -2.5 | 0.007* | 0.009* | 0.024 | 0.203 | 0.771 | (0.556, 0.986) |
| **493-5p** | 210 | 514.4 | -2.5 | 0.010* | 0.015 | 0.028 | 0.221 | 0.764 | (0.561, 0.967) |
| **543** | 223.1 | 753.2 | -3.3 | 2.5E-06* | 3.0E-04* | 2.0E-04* | 0.143 | 0.917 | (0.808, 1) |
| **654-3p** | 110.2 | 342.5 | -3.3 | 2.2E-04* | 0.009* | 0.006* | 0.193 | 0.826 | (0.642, 1) |
| **485-3p** | 93.2 | 352.3 | -3.3 | 5.8E-07* | 1.0E-04* | 3.3E-05* | 0.123 | 0.951 | (0.876, 1) |

*Abbreviations: CPM, miRNA counts per million; FC, fold change; error rates estimated by leave-one-out cross validation; AUROC, area under the receiver operating characteristic; CI, confidence interval; Significant Benjamini & Hochberg adjusted p-values are indicated by asterisks.*

**Figure 2.**
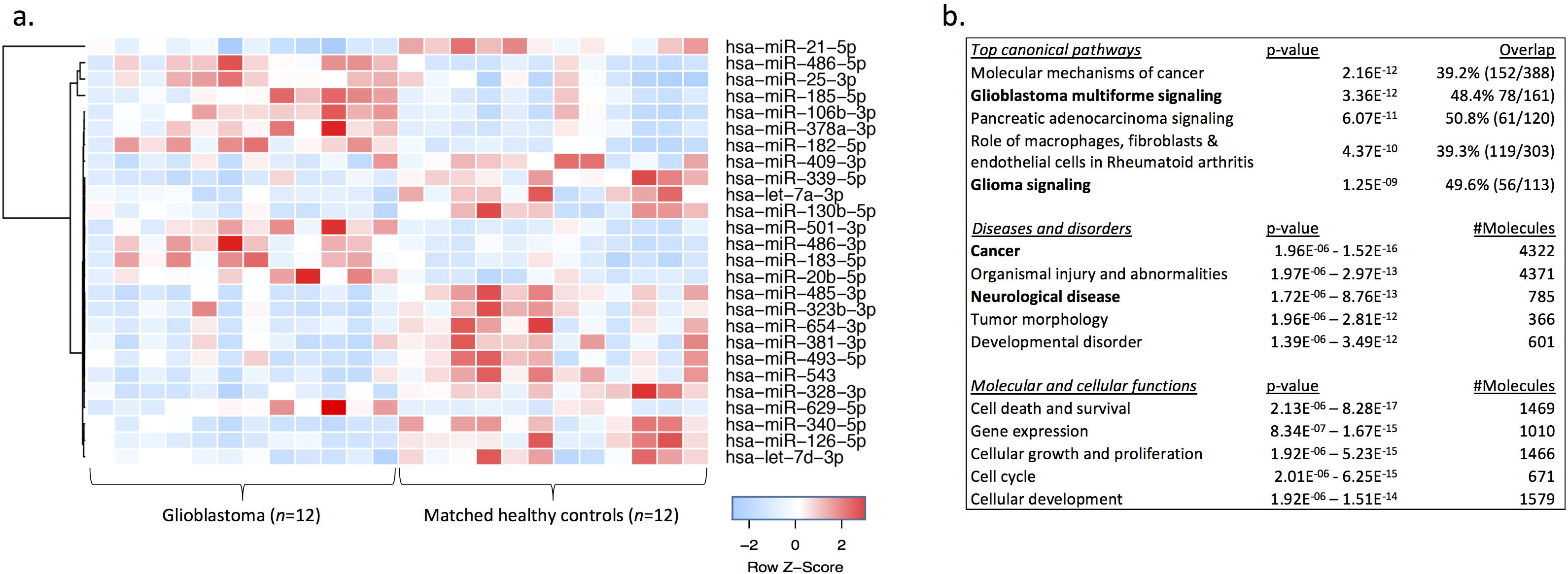
**(a.)** Hierarchical clustering of 26 differentially expressed miRNAs shows clear separation of glioblastoma (GBM) patients and healthy control (HC) exosomal profiles (fold change≥2 or ≤0.5; unadjusted *p*-values≤0.05 in all three statistical tests). **(b.)** Functional pathway analysis of mRNAs targeted by 44 significantly changing miRNA (unadjusted *p*-values≤0.05 in all three statistical tests) in GBM circulating exosomes. Top canonical pathways, diseases and disorders and molecular and cellular functions are listed with the numbers of overlapping molecules and significance of associations (right-tailed Fisher exact test, *p*-value).

*For more detailed demographic, clinical and histopathologic information, please refer to Supplementary Tables 1A-C. The mean age with standard deviation is provided for each cohort. Abbreviations: F, female; GBM, glioblastoma; GII-III, glioma grade II-III; GI_C, Ganglioglioma grade I control case; HC, healthy controls; M, male; MS, multiple sclerosis control cohort*

*Abbreviations: CPM, miRNA counts per million; FC, fold change; error rates estimated by leave-one-out cross validation; AUROC, area under the receiver operating characteristic; CI, confidence interval; Significant Benjamini & Hochberg adjusted p-values are indicated by asterisks.*

### Functional analysis of dysregulated miRNAs in GBM

We explored biological and canonical pathways associated with exosomal miRNAs changing in GBM patient sera relative to healthy controls. The identities of 44 miRNAs (*p*-value≤0.05 in all three tests; no fold change restriction) were uploaded into the IPA environment to analyze molecular pathways overrepresented in their targets. The dysregulated miRNAs target mRNAs that are significantly associated with ‘cancer’ (1.96E^-06^<*p*-value<1.52E^-16^) and ‘neurological disease’ (1.72E^-06^<*p*-value<8.76E^-13^) with around half of targeted mRNAs implicated in GBM (*p*-value=3.36E^-12^) and glioma signaling pathways (*p*-value=1.25E^-09^; **Fig. 2-b, Suppl.Fig.1**).

### Selection of signature miRNA classifiers for preoperative GBM diagnosis

The predictive power of each miRNA was estimated using LR models, in which individual miRNA expression profiles were used as predictors. ROC curves were determined and AUROC measures were ≥0.74 across the 26 dysregulated miRNAs. The 95% confidence intervals corresponding to AUROC estimates did not contain the null hypothesis value (AUROC=0.5 for a random prediction) indicating that all 26 miRNAs are statistically accurate univariate diagnostic predictors of GBM (**Table 2**; **Supp.Fig.2**). *In silico* validation by LOO-CV correctly identified the test sample on average 83% of the time (range 77–89%). We then used partitioning (70% training and 30% test) and Random Forest multivariate modeling to determine whether expression patterns of a subset of differentially expressed miRNAs could improve the predictive power. Using these methods, seven miRNAs (miR-182-5p, miR-328-3p, miR-339-5p, miR-340-5p, miR-485-3p, miR-486-5p and miR-543) distinguished GBM patients from healthy subjects in more than 75% of the random data partitions and were selected as the most ‘stable’ miRNA classifiers (**Fig.3a-b**). The RF model was repeated using all iterations of the seven most stable miRNAs and achieved an overall predictive power of 91.7% for classifying GBM patients from healthy controls (**Fig.3c**). The diagnostic accuracies of all possible combinations of the seven miRNAs were determined using AUROC measures along with the corresponding 95% confidence intervals (**Fig.3d; Supplementary Table 5)**. Strikingly, six miRNA combinations were able to distinguish GBM patients from healthy controls with perfect accuracy (**Fig. 3e**).

**Figure 3.**
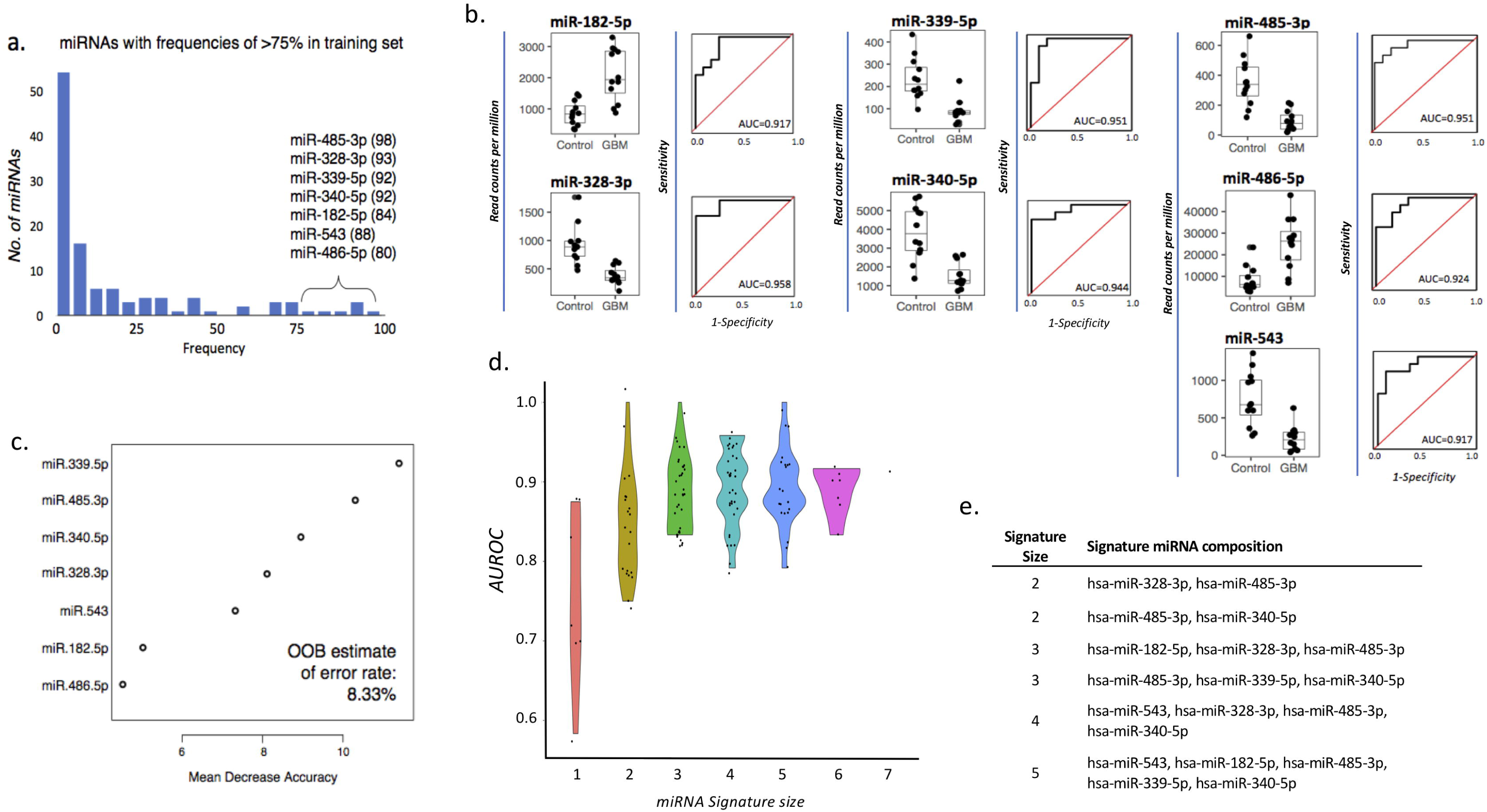
**(a.)** miRNAs appearing in >75 of 100 partitions (70% training set, 30% test set) were selected as the most stable miRNA classifiers by Random Forest modeling (frequencies are specified in brackets). **(b.)** Box-and-whisker plots and receiver operator characteristic curves with area under the curve (AUROC) calculations demonstrate the individual discriminatory power of the seven most stable miRNA classifiers. **(c.)** miRNAs were ordered by the importance of their contribution to discriminating GBM from [healthy] controls; overall out-of-the-bag (OOB) error rate of the seven features was 8.33%. **(d.)** AUROC measures of all possible combinations of the seven miRNAs previously identified to be the most stable predictors, stratified by the number of miRNAs (signature size) and their distributions and displayed as violin plots. **(e.)** miRNA signatures that discriminate between GBM and healthy controls with the perfect accuracy.

To assess the temporal stability of the GBM miRNA signature in the same patients, we tested preoperative sera collected at a GBM recurrence (GBM1 patient relapsed and required additional surgery after 8 months) and from an earlier GBM lesion (excised 4.6 months before GBM12; **Table 1B**). Using the panel of seven exosomal miRNAs, both GBM1-*relapse* and GBM12-*prior* were classified as GBM, in line with diagnostic histopathology. We also tested two independent samples, including a patient diagnosed with IDH^MUT^ GBM (GBM13) and a patient diagnosed with ‘high-grade glioma’ based on repeat MRIs and overall survival of 8.1 months (GBM14; see **Table 1B**). Both GBM13 and GBM14 were classified as GBM using the miRNA panel.

To further test the specificity of the GBM miRNA signature, we assessed its ability to distinguish GBM patients from additional healthy subjects and non-glioma disease controls. The panel accurately classified all additional healthy subjects (*n*=9; **Table 1B**) as well as a patient with ganglioglioma WHO (2016) grade I, a slow-growing, benign brain tumor with glioneuronal components (GIC-1). Next, we assessed the impact of neuroinflammatory disease processes on the specificity of our exosomal miRNA panel ability. The bioinformatics analysis above showed that dysregulated miRNAs also target mRNAs significantly associated with autoimmune rheumatoid arthritis and broadly to ‘neurological disease’ (**Fig. 2-b**). Our GBM miRNA panel was used to discriminate patients with the inflammatory autoimmune disease, multiple sclerosis (MS). Sera were sampled from MS patients with active gadolinium enhancing demyelinating lesions, either untreated or receiving immunomodulatory therapies (*n*=9; **Table 1B**). All MS patients were classified as controls, indicating the robustness of our exosomal miRNA signature for GBM identification.

### miRNAs dysregulated in IDH-mutant grade II-III gliomas provide additional markers for glioma severity and IDH mutational status

We then compared serum exosome miRNA profiles between IDH^MUT^ grade II-III glioma patients (*n*=10; mean age=42.7) and matched healthy controls (n=10; mean age=42.9; see **Table 1B**) and identified 23 differentially expressed miRNAs (fold change≥2; unadjusted *p*<0.05 in all three tests; **Supp.Table 4b.**). Of these, 12 miRNAs were shared with the GBM analysis and showed the same direction of change (**Fig. 4-a**). AUROC curve measures were ≥0.78 (average 0.88) across the 23 dysregulated miRNAs, and LOO-CV correctly identified the test sample on average 83% of the time (range 77–88%; **Supp.Table 5a.**; **Supp.Fig. 3a-b**). RF modeling performed on partitioned data selected miR-7d-3p, miR-98-5p, miR-106b-3p, 130b-5p and 185-5p as the most stable features for classifying grade II-III glioma patients from healthy participants, with a predictive power of 75.0% (**Fig. c-1.**; **Suppl.Fig.3c)**. The most stable miRNAs for classifying GII-III IDH^MUT^ from healthy controls were distinct from GBM IDH^WT^ signature miRNAs (**Fig.s 4b-1** and **4b-2**).

**Figure 4.**
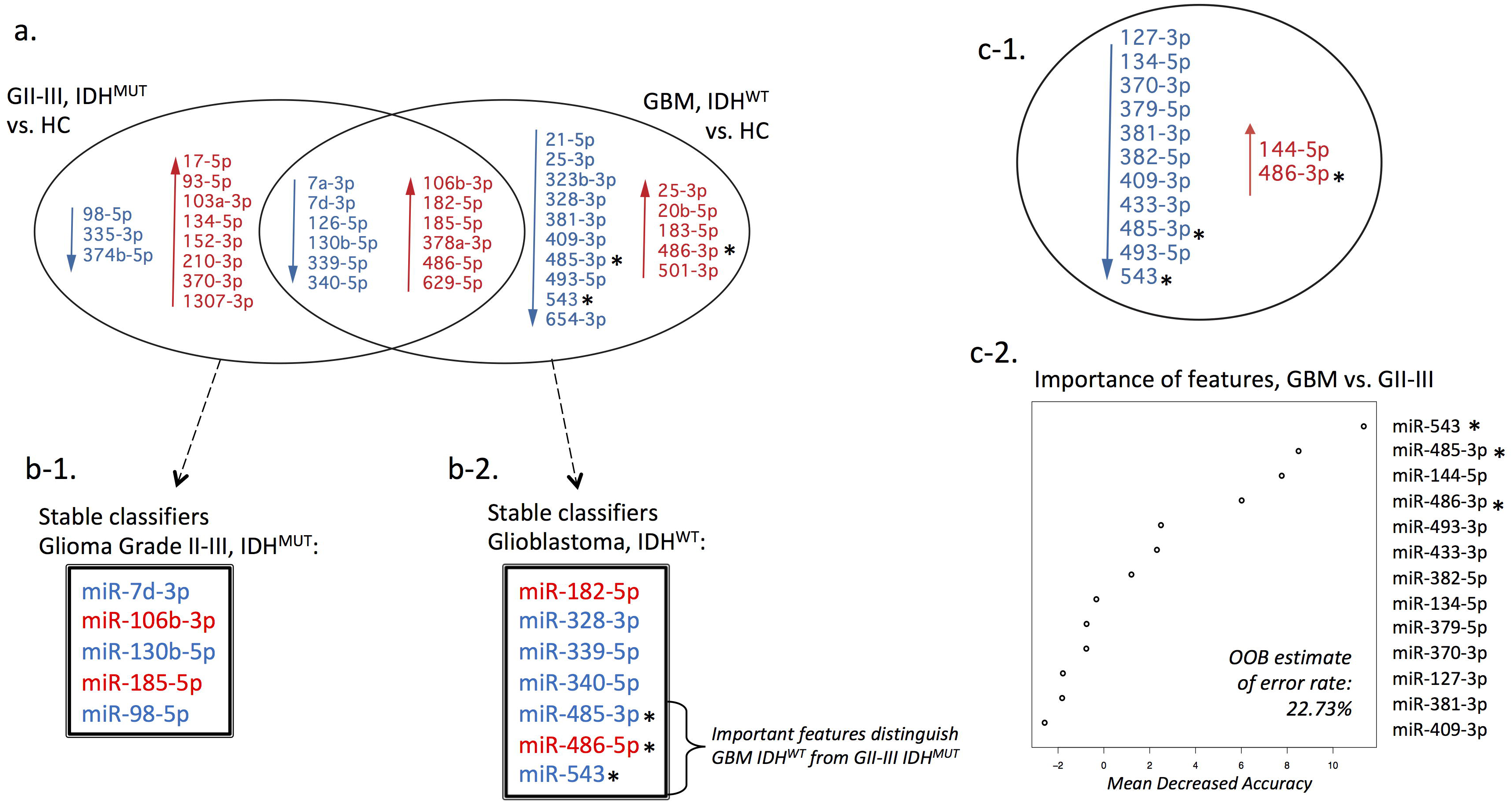
**(a.)** A Venn diagram summarizes the differentially expressed miRNAs between IDH^MUT^ glioma tumor grades II-III (GII-III; *n*=10), IDH^WT^ glioblastoma (GBM; *n*=12) and corresponding age- and gender-matched healthy controls (HC; fold change≥2 or ≤0.5; unadjusted *p*-values≤0.05 in all three statistics tests, i.e., Exact, *t*-test and Wilcoxon), with 12 overlapping differentially expressed miRNAs. Decreased expression is indicated in blue and increased expression in red. The most stable miRNAs for classifying (**b-1.**) GII-III IDH^MUT^ and (**b-2.**) GBM IDH^WT^ from HCs are listed and show distinct features. (**c-1.**) Summary of differentially expressed miRNAs between the GBM IDH^WT^ and GII-III IDH^MUT^ cohorts and (**c-2.**) plot of ‘importance’ of each individual miRNA for discriminating GBM from GII-III; out-of-the-bag (OOB) error rate is 22.73%. Three of the top four features that distinguish GBM IDH^WT^ from GII-III IDH^MUT^ were only identified in the GBM vs. HC comparative analysis, are members of the GBM miRNA signature that together accurately classify GBMs from HCs and may be specific markers for GBM (indicated by **asterisks in a.**, **b-2., c-1., and c-2.**).

The sncRNA data was further interrogated to ascertain whether a subset of miRNAs showed potential for distinguishing glioma disease severity or IDH mutational status. Direct comparisons between GBM IDH^WT^ and GII-III IDH^MUT^ patients revealed 13 differentially expressed miRNAs (fold change≥2; unadjusted *p*<0.05 in all three tests; (**Fig. 4c-1.; Supp.Table 4c**). AUROC curve measurements were ≥0.78 (average 0.84) across the 13 dysregulated miRNAs and LOO-CV correctly identified the test sample on average 80% of the time (range 76–86%; **Supp.Table 5b.**; **Supp.Fig. 4a-b**). Numbers of significant miRNA were too few to perform partitioning, so a single RF model was constructed from all 13 dysregulated miRNAs that showed an estimated predictive power of 77.4% (**Fig. 4c-2.**) Interestingly, three of the top four features that discriminate GBM IDH^WT^ from GII-III IDH^MUT^ are members of the GBM miRNA signature (i.e., miR-543, miR-485-3p and miR-486-3p), changing only in GBM patient sera relative to healthy participants (indicated by asterisks in **Fig. 4)**.

## DISCUSSION

Using unbiased high-throughput next generation sequencing and an integrative bioinformatics pipeline^8^, we have identified differentially expressed serum exosomal miRNAs that discriminate GBM patients from healthy controls. Machine-learning approaches on miRNAs were used to examine their individual and shared predictive abilities for a pre-operative GBM diagnosis via a blood test. Of the 26 differentially expressed miRNAs in GBM patients’ relative to healthy controls, we selected a stable signature panel of seven miRNAs. Together, expression levels of miR-182-5p, miR-328-3p, miR-339-5p, miR-340-5p, miR-485-3p, miR-486-5p and miR-543 predicted a preoperative GBM diagnosis with a 91.7% accuracy. Within this multivariate model a combination of just four miRNAs (miR-182-5p, miR-328-3p miR-485-3p miR-486-5p) distinguished GBM patients from healthy controls with perfect accuracy (100.0%).

There have been multiple studies examining ‘free-circulating’ miRNAs in glioma patients with varying success. A recent meta-analysis of these studies found the specificity and sensitivity of circulating miRNAs was 0.87 and 0.86, respectively, while noting the large heterogeneity of circulating miRNAs within the included studies ^10^. The heterogeneity is likely due to differences in data normalization used in qRT-PCR studies, with no universally accepted endogenous housekeeping control ^10^. Interestingly, the majority of miRNAs identified in our exosomal signature have not been previously identified in ‘free-circulating’ studies. This is consistent with the notion that exosomes represent a distinct pathway of nucleic acid release from cells, and contain selectively packaged miRNA species ^5^. We have previously shown the effects of RNAse pre-treatment of serum prior to exosome isolation, as performed in this study, drastically alters the miRNA profiles identified, presumably due to eradication of co-precipitated ‘free-circulating’ miRNAs ^8^. Moreover, normalization of deep sequencing data is not dependant on comparison to a reference signal or housekeeping gene, potentially reducing variability in data analysis.

Functional pathway analysis of mRNA species targeted by exosomal miRNAs dysregulated in GBM patient sera showed highly significant associations to specific GBM molecular pathways. This provides confidence that the miRNA biomarkers resolved by our methods are relevant to this particular disease setting. Previous studies have identified roles for all seven GBM miRNA classifiers in various aspects of glioma and GBM biology. miR-182, detected here in significantly higher levels in GBM sera, was proposed as a marker of glioma progression, critical for glioma tumorigenesis, tumor growth and survival *in vitro*^11^,^12^, with high miR-182 tissue expression observed in GBM^13^ and associated with poor overall survival^14^. Also in line with observations here, the up-regulation of miR-486 was shown to promote glioma aggressiveness both *in vitro* and *in vivo*^15^. Exosomal miRNAs identified with lower expression levels in GBM patient sera are also substantiated by the literature. Functional assays indicate tumor suppressive roles of miR-328^16^, miR-340^17^,^18^, miRNA-485-5p^19^ and miR-543^20^ with low levels observed in tumor tissues relative to normal brain^16^,^18^-^20^ and low tissue expression levels significantly associated to poor patient outcomes^16^,^18^. While miR-339 (decreased levels in GBM patients here) was shown to contribute to immune evasion of GBM cells by modulating T-cell responses^21^, inhibitory roles for miR-339 were reported in acute myeloid leukemia^22^, hepatocellular carcinoma^23^, gastric^24^, colorectal^25^, breast^26^ and ovarian cancers^27^.

The GBM miRNA signature was able to accurately classify all additional specimens in the validation sets (healthy, *n*=9; non-glioma, *n*=10), including patients with gadolinium enhancing active demyelinating lesions. Tumefactive demyelination is a well-recognized mimic of GBM^28^. The GBM signature also correctly classified four additional GBM specimens, including two serial collections from patients within the discovery cohort as well as two independent patients. This pilot study utilised a relatively small patient group, and further testing is needed to determine whether the miRNA panel can reliably diagnose GBM in large, independent patient cohorts. Moreover, the correlation between a positive GBM classification and tumor burden needs to be addressed. To this end, longitudinal studies should be pursued to assess whether the GBM miRNA panel can detect time critical GBM tumor recurrences.

There is more than one pathological route to a GBM; primary and secondary GBMs are distinct entities with IDH mutations considered a genetic signpost ^29^. The only patients where early detection of a GBM tumor is likely are arguably those with diffuse and anaplastic (grade II-III) gliomas who progress with a secondary GBM recurrence (IDH^MUT^). Accordingly, the identification of reliable and readily accessible circulating progression markers is an important step towards precision medicine for patients diagnosed with low grade gliomas. While the GBM miRNA signature was described in serum exosomes from IDH^WT^ GBM patients, it was also able to categorize a patient with IDH^MUT^ GBM (GBM13) from healthy participants. It is worth noting that miRNA members of the GBM signature panel (specifically, increased miR-182-5p, decreased miR339-5p and miR-340-5p) were also identified in the IDH^MUT^ GII-III comparative analysis. Whether these miRNA changes are related to IDH mutational status, glioma grade, or a combination of the two, cannot be delineated here. However, our multivariate modeling did identify distinct panels of miRNAs for classifying GBM and glioma patients from their corresponding matched healthy control cohorts. Moreover, three GBM signature panel miRNAs that were unique to the GBM vs control comparative analysis (increased miR-486-5p and decreased miR-485-3p and miR-543) were among the top four features that distinguish GBM IDH^WT^ from GII-III IDH^MUT^ and therefore, might be specific for GBM IDH^WT^ (indicated by asterisks in **Fig.4**). These encouraging results demonstrate the potential for exosomal miRNA profiles to be used for glioma subtyping and grading, including the determination of mutational states. Expansion of these discovery analyses to include well defined cohorts of glioma subtypes with sufficient *n*, will likely resolve biomarkers of more nuanced specificity.

## SUMMARY

In summary, we have described a serum exosomal miRNA signature that can accurately predict a GBM diagnosis, preoperatively. This pilot study demonstrates that exosomal associated miRNAs have exceptional utility as biomarkers in the glioma disease setting. If these exosomal biomarkers are able to offer non-invasive, early indications of tumor progression and/or recurrence, they are likely to have significant clinical utility. These exciting findings have significant potential to transform current diagnostic paradigms, as well as provide distinct surrogate endpoints for clinical trials. Assessment of serum exosomal miRNAs in larger longitudinal cohorts of patients with GBM are required to definitely determine their utility in clinical practice, and these studies are currently underway.

## METHODS

### Participants

Serum (1 mL) was accessed from the Neuropathology Tumor and Tissue Bank at Royal Prince Alfred Hospital, New South Wales, Australia (Sydney Local Health District HREC approval, X014-0126 & HREC/09RPAH/627). Twenty-six serum specimens were collected pre-operatively from patients with histologically confirmed glioma tumors, including 16 with GBM, IDH-wildtype (IDH^WT^) WHO (2016) grade IV, and 10 patients with grade II-III IDH-mutant (IDH^MUT^) gliomas (refer to **Table 1**; **Supp.Table 1** for more detailed information). Age- and gender-matched healthy control sera (*n*=16) were used for discovery miRNA analyses. Sera from an additional nine healthy controls and ten non-glioma patients (including active MS, *n*=9, and ganglioglioma, *n*=1) were used to test the GBM miRNA signature. This study was performed under RPAH, and USYD HREC approved protocols (#X13-0264 and 2012/1684), and all participants provided written informed consent. All methods were performed in accordance with the relevant guidelines and regulations.

### Exosome purification and characterization

Exosomes were isolated from serum as previously described^8^. Briefly, serum (1 mL from each subject) was treated with RNase A (37☐°C for 10☐min; 100 ng/ml; Qiagen, Australia) before exosome purification by size exclusion chromatography (qEV iZONE Science). Ten fractions (500☐μL) were eluted in PBS, as per manufacturer’s instructions. Fractions 8, 9, and 10 were previously shown to contain purified exosome populations^8^ and were collected and stored at −80☐°C. Captured exosomes were characterized in accordance with the criteria outlined by the International Society for Extracellular Vesicles (ISEV)^30^. Specifically, we identified more than three exosome-enriched proteins by mass spectrometry proteome profiling and characterized vesicle heterogeneity using two technologies, transmission electron microscopy (TEM) and nanoparticle tracking analysis (NTA).

#### Transmission electron microscopy

Combined qEV-captured fractions 8-10 were loaded onto carbon-coated, 200 mesh Cu formvar grids (#GSCU200C; ProSciTech Pty Ltd, QLD, Australia), fixed (2.5% glutaraldehyde, 0.1 M phosphate buffer, pH7.4), negatively stained with 2% uranyl acetate for 2☐min and dried overnight. Exosomes were visualised at 40,000 X magnification on a Philips CM10 Biofilter TEM (FEI Company, OR, USA) equipped with an AMT camera system (Advanced Microscopy Techniques, Corp., MA, USA) at an acceleration voltage of 80☐kV.

#### Nanoparticle tracking analysis

Particle size distributions and concentrations were measured by NTA software (version 3.0) using the NanoSight LM10-HS (NanoSight Ltd, Amesbury, UK), configured with a 532-nm laser and a digital camera (SCMOS Trigger Camera). Video recordings (60 s) were captured in triplicate at 25 frames/s with default minimal expected particle size, minimum track length, and blur setting, a camera level of 10 and detection threshold of 5.

#### Proteome analysis of exosomal preparations

Serum exosome fractions 8, 9 and 10 were prepared for mass spectrometry (MS)-based proteomic analysis. Proteomes were concentrated using chloroform-methanol precipitation, dissolved in 90% formic acid (FA), their concentrations estimated at 280 nm using a Nanodrop (ND-1000, Thermo Scientific, USA) and aliquots dried using vacuum centrifugation. Proteomes were then processed and quantified as before ^31^. Peptides from each fraction were desalted using C18 ZipTips™, concentrations estimated by Qubit quantitation (Invitrogen), dried by vacuum centrifugation and re-suspended in 3% acetonitrile (ACN; v/v)/0.1% formic acid (v/v). Samples (0.5 μg) from exosome elution fractions 8-10 were separated by nanoLC using an Ultimate nanoRSLC UPLC and autosampler system (Dionex) before analyzed on a QExactive Plus mass spectrometer (Thermo Electron, Bremen, Germany) as previously described^31^. MS/MS data were analyzed using Mascot (Matrix Science, London, UK; v2.4.0) with a fragment ion mass tolerance of 0.1 Da and a parent ion tolerance of 4.0 PPM. Peak lists were searched against a SwissProt database (2017_11), selected for *Homo sapiens*, trypsin digestion, max. 2 missed cleavages, and variable modifications methionine oxidation and cysteine carbamidomethylation. Exosome proteins were annotated using Vesiclepedia (http://microvesicles.org)^32^ and Functional Enrichment Analysis Tool (FunRich; v2.1.2; http://funrich.org)^33^.

### RNA extraction and small RNA sequencing

Serum exosomes were processed for RNA extraction using the Plasma/Serum Circulating & Exosomal RNA Purification Mini Kit (Norgen Biotek, Cat. 51000) according to the manufacturer’s protocol. Extracted total RNA samples were analyzed with a Eukaryote Total RNA chip on an Agilent 2100 Bioanalyser (Agilent Technologies, United States) to confirm sufficient yield, quality and size of RNA. Exosome RNA sequencing libraries were then constructed using the NEBNext Multiplex Small RNA Library Prep Kit for Illumina (BioLabs, New England) according to the manufacturer’s instructions. Yield and size distribution of resultant libraries were validated using Agilent 2100 Bioanalyzer on a High-sensitivity DNA Assay (Agilent Technologies, United States). Libraries were then pooled with an equal proportion for multiplexed sequencing on Illumina HiSeq. 2000 System at the Ramaciotti Centre for Genomics.

### Data pre-processing, differential expression analysis and pathway analysis

Data pre-processing was performed using a pipeline comprising of adapter trimming (cutadapt), followed by genome alignment to human genome hg 19 using Bowtie (18☐bp seed, 1 error in seed, quality score sum of mismatches<70). Where multiple best strata alignments existed, tags were randomly assigned to one of those coordinates. Tags were annotated against mirBase 20 and filtered for at most one base error within the tag. Counts for each miRNA were tabulated and adjusted to counts per million miRNAs passing the mismatch filter. All samples achieved miRNA read counts >45,000 read counts and miRNAs with low abundance (<50 read counts across more than 20% of samples) were removed. Differential expression analysis was performed using three different statistical hypothesis tests including a non-parametric two-sample Wilcoxon test and two parametric tests-Student’s *t*-test, and an Exact test (implemented in Bioconductor EdgeR), which tests for differences between the means of two groups of negative-binomially distributed counts. Benjamini & Hochberg adjusted *p*-values were also calculated. Data pre-processing and differential expression analysis were performed using Bioconductor and R statistical packages. Pathway analysis was performed using Ingenuity® software (Ingenuity Systems, USA; http://analysis.ingenuity.com). MicroRNA target filters were applied to significant, differentially expressed miRNAs (unadjusted *p*-value≤0.05 in all three statistical methods) and mRNA target lists were generated based on highly predicted or experimentally observed confidence levels. Core expression analyses were performed with default criteria to determine the most significant functional associations (biological and canonical pathways) of mRNAs targeted by dysregulated miRNAs.

### Univariate analysis

We performed logistic regression (LR) and receiver operator characteristic (ROC) analysis to assess the predictive power of individual miRNAs between the two groups of interest. LR was used to identify linear predictive models with each miRNA as the univariate predictor. The quality of each model was depicted by the corresponding ROC curve, which plots the true positive rate (i.e., sensitivity) against the false-positive rate (i.e., 1-specificity). The area under the ROC curve (AUROC) was then computed as a measure of how well each LR model can distinguish between two diagnostic groups. The 95% confidence intervals (CI) of AUROC measures were estimated using Delong method^34^ to assess the significance of a model’s predictive power as compared to a random trial (i.e., AUROC = 0.5). We then used leave-one-out cross-validation (LOO-CV) to estimate the prediction errors of the LR models. LOO-CV learns the model on all samples except one and tests the learnt model on the left-out sample. The process is repeated for each sample and the error rate is the proportion of misclassified samples. Overall, cross validation is a powerful model validation technique for assessing how the results of a statistical analysis can be generalized to an independent dataset^35^. These analyses were performed using R stats (glm) and boot (cv.glm) packages.

### Multivariate Analysis

To assess the predictive power of multiple miRNAs as disease signatures, samples were first randomly partitioned into two disjoint sets of *discovery* (70% of samples) and *validation* (30% of samples). MiRNAs differentially expressed in the discovery set (i.e., changes increased or decreased by fold change≥2 and unadjusted *p*-value≤0.05 in all three statistical hypothesis tests) were then selected as features/predictors of *Random Forest* (RF) multivariate predictive model. RF is a multivariate nonlinear classifier that operates by constructing a multitude of decision trees at training time in order to correct for the overfitting problem^36^. RF was trained on the *discovery* set and the resultant predictive model was then used to predict GBM or GII-III patients *versus* healthy controls based on the read count values of identified miRNAs in *validation* samples. For statistical rigour, to account for random partitioning of the samples into discovery and validation sets, the whole process was repeated 100 times. We then chose *stable* miRNAs—i.e., those identified to be differentially expressed in more than 75% of iterations—as predictors of an RF model using all samples and the AUROC with 95% CI as well as out-of-bag (OOB) error was reported as an unbiased estimates of the model predictive power. The ‘*importance*’ or relative contribution of each feature (differentially expressed miRNAs) in the RF performance was then estimated based on the ‘*mean decrease accuracy*’ measure as discussed in^37^. All analyses were performed using R ‘caret’ and ‘RandomForest’ packages.

### Data Availability

Exosomal miRNA raw data *will be* accessible at NCBI Gene Expression Omnibus (GEO; *accession number to be provided*). In the interim, the miRNA sequencing data is available at: https://github.com/VafaeeLab/glioblastoma_exosomal_miR_markers. Normalised data used for statistical analysis is provided in Supplementary Table 3.

## Acknowledgements

Many thanks to the wonderful and dedicated staff at RPAH, with special thanks to Mary Lordan, Jane Raftesath, Audrey Caudon, Shu Wang and Ladan Noroozi.

## Contributions

All authors contributed to manuscript preparation and approve the submission of the work presented here. Specific contributions are as follows: S.E. performed technical work including serum processing, exosome purification, electron microscopy, small RNA sequencing, data analysis, manuscript preparation. F.V. and P.Y. developed and performed the bioinformatics analytical pipeline. S.H. performed technical work including serum processing, nanosight particle tracking, and mass spectrometry. H.W and M.Y.T.L characterized clinical cases, including molecular characterizations of tumour tissue. L.S and B.S assisted with clinical sample procurement and case characterization. C.M.S. assisted with small RNA sequencing protocols and data interpretation. M.E.B. provided experimental design and data interpretation. K.L.K provided experimental design, cohort characterisation, proteomics methods, bioinformatics, data interpretation and presentation. M.E.B and K.L.K are guarantors of this work.

## REFERENCES

1 Ellingson, B. M., Wen, P. Y. & Cloughesy, T. F. Modified Criteria for Radiographic Response Assessment in Glioblastoma Clinical Trials. Neurotherapeutics: the journal of the American Society for Experimental NeuroTherapeutics 14, 307–320, doi:10.1007/s13311-016-0507-6 (2017).

2 Saadatpour, L. et al. Glioblastoma: exosome and microRNA as novel diagnosis biomarkers. Cancer gene therapy 23, 415–418, doi:10.1038/cgt.2016.48 (2016).

3 Skog, J. et al. Glioblastoma microvesicles transport RNA and proteins that promote tumour growth and provide diagnostic biomarkers. Nature cell biology 10, 1470–1476, doi:10.1038/ncb1800 (2008).

4 Mallawaaratchy, D. M. et al. Comprehensive proteome profiling of glioblastoma-derived extracellular vesicles identifies markers for more aggressive disease. J Neurooncol 131, 233–244, doi:10.1007/s11060-016-2298-3 (2017).

5 Li, C. C. et al. Glioma microvesicles carry selectively packaged coding and non-coding RNAs which alter gene expression in recipient cells. RNA biology 10, 1333–1344, doi:10.4161/rna.25281 (2013).

6 Akers, J. C. et al. MiR-21 in the extracellular vesicles (EVs) of cerebrospinal fluid (CSF): a platform for glioblastoma biomarker development. PloS one 8, e78115, doi:10.1371/journal.pone.0078115 (2013).

7 Manterola, L. et al. A small noncoding RNA signature found in exosomes of GBM patient serum as a diagnostic tool. Neuro-oncology 16, 520–527, doi:10.1093/neuonc/not218 (2014).

8 Ebrahimkhani, S. et al. Exosomal microRNA signatures in multiple sclerosis reflect disease status. Scientific reports 7, 14293, doi:10.1038/s41598-017-14301-3 (2017).

9 Roberts, T. C. The MicroRNA Biology of the Mammalian Nucleus. Molecular therapy. Nucleic acids 3, e188, doi:10.1038/mtna.2014.40 (2014).

10 Ma, C. et al. A comprehensive meta-analysis of circulation miRNAs in glioma as potential diagnostic biomarker. PLoS One 13, e0189452, doi:10.1371/journal.pone.0189452 (2018).

11 Hailin, T. et al. The miR-183/96/182 Cluster Regulates Oxidative Apoptosis and Sensitizes Cells to Chemotherapy in Gliomas. Current Cancer Drug Targets 13, 221–231, doi:http://dx.doi.org/10.2174/1568009611313020010 (2013).

12 Xue, J. et al. miR-182-5p Induced by STAT3 Activation Promotes Glioma Tumorigenesis. Cancer research 76, 4293–4304, doi:10.1158/0008-5472.CAN-15-3073 (2016).

13 Kouri, F. M. et al. miR-182 integrates apoptosis, growth, and differentiation programs in glioblastoma. Genes & Development 29, 732–745, doi:10.1101/gad.257394.114 (2015).

14 Jiang, L. et al. miR-182 as a Prognostic Marker for Glioma Progression and Patient Survival. The American Journal of Pathology 177, 29–38, doi:10.2353/ajpath.2010.090812 (2010).

15 Song, L. et al. miR-486 sustains NF-κB activity by disrupting multiple NF-?B-negative feedback loops. Cell Research 23, 274–289, doi:10.1038/cr.2012.174 (2013).

16 Yuan, J. et al. microRNA-328 is a favorable prognostic marker in human glioma via suppressing invasive and proliferative phenotypes of malignant cells. International Journal of Neuroscience 126, 145–153, doi:10.3109/00207454.2014.1002610 (2016).

17 Li, X. et al. miR-340 inhibits glioblastoma cell proliferation by suppressing CDK6, cyclin-D1 and cyclin-D2. Biochemical and Biophysical Research Communications 460, 670–677, doi:https://doi.org/10.1016/j.bbrc.2015.03.088 (2015).

18 Huang, D. et al. miR-340 suppresses glioblastoma multiforme. Oncotarget 6, 9257–9270 (2015).

19 Yu, J., Wu, S.-W. & Wu, W.-P. A tumor-suppressive microRNA, miRNA-485-5p, inhibits glioma cell proliferation and invasion by down-regulating TPD52L2. American Journal of Translational Research 9, 3336–3344 (2017).

20 Xu, L. et al. miR-543 functions as a tumor suppressor in glioma in vitro and in vivo. Oncology Reports 38, 725–734, doi:10.3892/or.2017.5712 (2017).

21 Ueda, R. et al. Dicer-regulated microRNAs 222 and 339 promote resistance of cancer cells to cytotoxic T-lymphocytes by down-regulation of ICAM-1. Proceedings of the National Academy of Sciences of the United States of America 106, 10746–10751, doi:10.1073/pnas.0811817106 (2009).

22 Barrera-Ramirez, J. et al. Micro-RNA Profiling of Exosomes from Marrow-Derived Mesenchymal Stromal Cells in Patients with Acute Myeloid Leukemia: Implications in Leukemogenesis. Stem Cell Reviews 13, 817–825, doi:10.1007/s12015-017-9762-0 (2017).

23 Wang, Y.-L., Chen, C.-m., Wang, X.-M. & Wang, L. Effects of miR-339-5p on invasion and prognosis of hepatocellular carcinoma. Clinics and Research in Hepatology and Gastroenterology 40, 51–56, doi:https://doi.org/10.1016/j.clinre.2015.05.022 (2016).

24 Shen, B. et al. MicroRNA-339, an epigenetic modulating target is involved in human gastric carcinogenesis through targeting NOVA1. FEBS Letters 589, 3205–3211, doi:10.1016/j.febslet.2015.09.009 (2015).

25 Zhou, C., Lu, Y. & Li, X. miR-339-3p inhibits proliferation and metastasis of colorectal cancer. Oncology Letters 10, 2842–2848, doi:10.3892/ol.2015.3661 (2015).

26 Wu, Z.-s. et al. MiR-339-5p inhibits breast cancer cell migration and invasion in vitro and may be a potential biomarker for breast cancer prognosis. BMC Cancer 10, 542–542, doi:10.1186/1471-2407-10-542 (2010).

27 Shan, W., Li, J., Bai, Y. & Lu, X. miR-339-5p inhibits migration and invasion in ovarian cancer cell lines by targeting NACC1 and BCL6. Tumor Biology 37, 5203–5211, doi:10.1007/s13277-015-4390-2 (2016).

28 Riva, D. et al. A case of pediatric tumefactive demyelinating lesion misdiagnosed and treated as glioblastoma. J Child Neurol 23, 944–947, doi:10.1177/0883073808315419 (2008).

29 Ohgaki, H. & Kleihues, P. The Definition of Primary and Secondary Glioblastoma. Clinical Cancer Research 19, 764–772, doi:10.1158/1078-0432.Ccr-12-3002 (2013).

30 Lotvall, J. et al. Minimal experimental requirements for definition of extracellular vesicles and their functions: a position statement from the International Society for Extracellular Vesicles. Journal of extracellular vesicles 3, 26913, doi:10.3402/jev.v3.26913 (2014).

31 Mallawaaratchy, D. M. et al. Membrane proteome analysis of glioblastoma cell invasion. J Neuropathol Exp Neurol 74, 425–441, doi:10.1097/NEN.0000000000000187 (2015).

32 Kalra, H. et al. Vesiclepedia: a compendium for extracellular vesicles with continuous community annotation. PLoS biology 10, e1001450, doi:10.1371/journal.pbio.1001450 (2012).

33 Pathan, M. et al. FunRich: An open access standalone functional enrichment and interaction network analysis tool. Proteomics, doi:10.1002/pmic.201400515 (2015).

34 DeLong, E. R., DeLong, D. M. & Clarke-Pearson, D. L. Comparing the areas under two or more correlated receiver operating characteristic curves: a nonparametric approach. Biometrics 44, 837–845 (1988).

35 Seni, G. & Elder, J. F. Ensemble Methods in Data Mining: Improving Accuracy Through Combining Predictions. Synthesis Lectures on Data Mining and Knowledge Discovery 2, 1–126, doi:10.2200/s00240ed1v01y200912dmk002 (2010).

36 Hastie, T., Robert, T. & Friedman, J. Elements of Statistical Learning: data mining, inference, and prediction. (Springer, 2008).

37 Breiman, L. Random Forests. Machine Learning 45, 5–32, doi:10.1023/A:1010933404324 (2001).

